# Salvage of Ribose from Uridine or RNA Supports Glycolysis when Glucose is Limiting

**DOI:** 10.1101/2021.06.10.447789

**Authors:** Alexis A. Jourdain, Owen S. Skinner, Akinori Kawakami, Russel P. Goodman, Hongying Shen, Lajos V. Kemény, Lena Joesch-Cohen, Matthew G. Rees, Jennifer A. Roth, David E. Fisher, Vamsi K. Mootha

## Abstract

Glucose is vital for life, serving both as a source of energy and as a carbon building block for growth. When glucose availability is limiting, alternative nutrients must be harnessed. To identify mechanisms by which cells can tolerate complete loss of glucose, we performed nutrient-sensitized, genome-wide genetic screening and growth assays of 482 pooled PRISM cancer cell lines. We report that catabolism of uridine enables the growth of cells in the complete absence of glucose. While previous studies have shown that the uracil base of uridine can be salvaged to support growth in the setting of mitochondrial electron transport chain deficiency (*1*), our work shows that the ribose moiety of uridine can be salvaged via a pathway we call “uridinolysis” defined as: [1] the phosphorylytic cleavage of uridine by UPP1/2 into uracil and ribose-1-phosphate (R1P), [2] the conversion of R1P into fructose-6-P and glyceraldehyde-3-P by PGM2 and the non-oxidative branch of the pentose phosphate pathway (non-oxPPP), and [3] their glycolytic utilization to fuel ATP production, biosynthesis and gluconeogenesis. Intriguingly, we report that uridine nucleosides derived from RNA are also a substrate for uridinolysis and that RNA can support growth in glucose-limited conditions. Our results underscore the malleability of central carbon metabolism and raise the provocative hypothesis that RNA can also serve as a potential storage for energy.

## Main Text

To identify novel genes and pathways involved in energy metabolism, we transduced K562 cells with a library comprising 17,254 barcoded ORFs (*2*) and compared proliferation in media containing glucose and galactose, a poor substrate for glycolysis. We used DMEM that contained glutamine, as well as pyruvate and uridine, for which cells with OXPHOS defects are dependent (*1*). After 21 days, we harvested cells and sequenced their barcodes using next-generation sequencing (Table S1). The most striking result was a strong enrichment in galactose for ORFs encoding *UPP1* and *UPP2*, which encode two paralogous uridine phosphorylases catalyzing the phosphate-dependent catabolism of uridine into R1P and uracil (Fig. 1B, S1). To confirm our screening result, we stably expressed *UPP1* and *UPP2* ORFs in K562 cells and observed a significant gain in proliferation in galactose media (Fig. 1C, S1C). This gain was dependent on uridine being present in the media, while expression of *UPP1/2*, or addition of uridine, had no effects in glucose-containing media. Importantly, we found that *UPP1*-expressing cells also efficiently proliferated in media containing uridine in the complete absence of glucose or galactose (“sugar-free”), while control cells were unable to proliferate (Fig. 1D). The ability of *UPP1* cells to grow in the sugar-free media strictly depended on uridine, which could not be substituted by any of the other seven nucleosides precursors of nucleic acids (Fig. 1E). Furthermore, expression of *UPP1* alone was sufficient to enable growth on sugar-free media supplemented with purified RNA (Fig. 1F). We conclude that a strong uridine phosphorylase activity confers the ability to grow in media containing uridine or RNA, in the complete absence of glucose.

**Figure 1:**
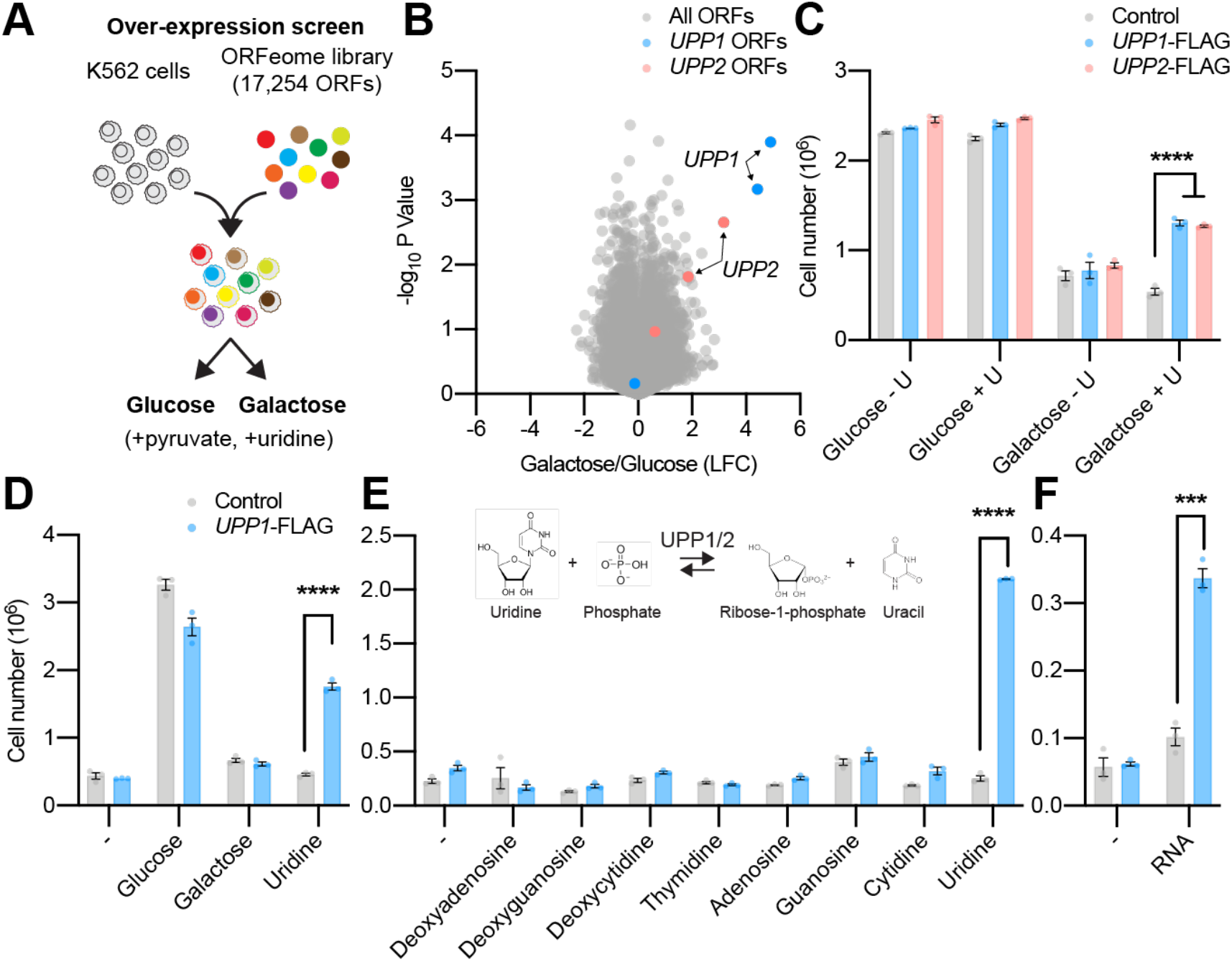
Uridine Phosphorylase Activity Supports Growth on Uridine and RNA. (A) Schematic overview of the ORF proliferation screen. (B) Volcano plot representation of the screen hits after 21 days of growth in media containing 25mM of glucose or 25mM galactose, 0.2mM uridine and 1mM sodium pyruvate. (C) Cell growth assays of K562 cells expressing *GFP* or *UPP1*-FLAG in pyruvate-free media containing 25mM of glucose or 25mM of galactose, in the presence or absence of 0.2mM uridine (+/− U), (D) 10mM of either glucose, galactose or uridine, (E) 5mM of the indicated nucleosides, or (F) 0.5mg/mL of purified yeast RNA. All data are shown as mean ±SEM with ***p<10^−3^ and ****p<10^−4^ t-test relative to control, with *n*=3 or 4.

We next took a complementary approach and asked in a systematic way whether a diverse collection of human cancer cell lines have differential latent ability to grow on uridine or RNA, when glucose is absent. We screened 482 pooled barcoded adherent cancer cell lines comprising 22 different lineages from the PRISM collection (*3*) in media containing 10mM glucose or uridine, in the absence of any supplemental sugar (Fig. 2A). Cells from the melanoma and the glioma lineages grew remarkably well in uridine as compared to the other lineages, whereas Ewing sarcoma cells grew significantly less (Fig. 2B). Cell lines from the PRISM collection have been extensively characterized at a molecular level (*4*), so we searched for baseline genomic factors that correlate with the ability to grow on uridine (Table S2). Consistent with our ORF screen, we found that across the 482 cell lines the ability to grow on uridine correlated best with *UPP1* expression, both at the transcript and the protein levels, as well as in the gene copy number space (Fig. 2C-E). Expression of *UPP1* across the PRISM collection was the highest in cell lines of skin origin (Fig. S2A), where high uridine phosphorylase enzyme activity has been documented (*5*). In agreement with this, we found significant, *UPP1*-dependent, proliferation of melanoma cells in sugar-free media supplemented with uridine or RNA (Fig. 2F, G). The correlation between *UPP1* expression and growth on uridine extended to other lineages as well (Fig. S2B-C), while *UPP2* was almost never expressed in the PRISM collection (TPM<1) (Fig. S2A).

**Figure 2:**
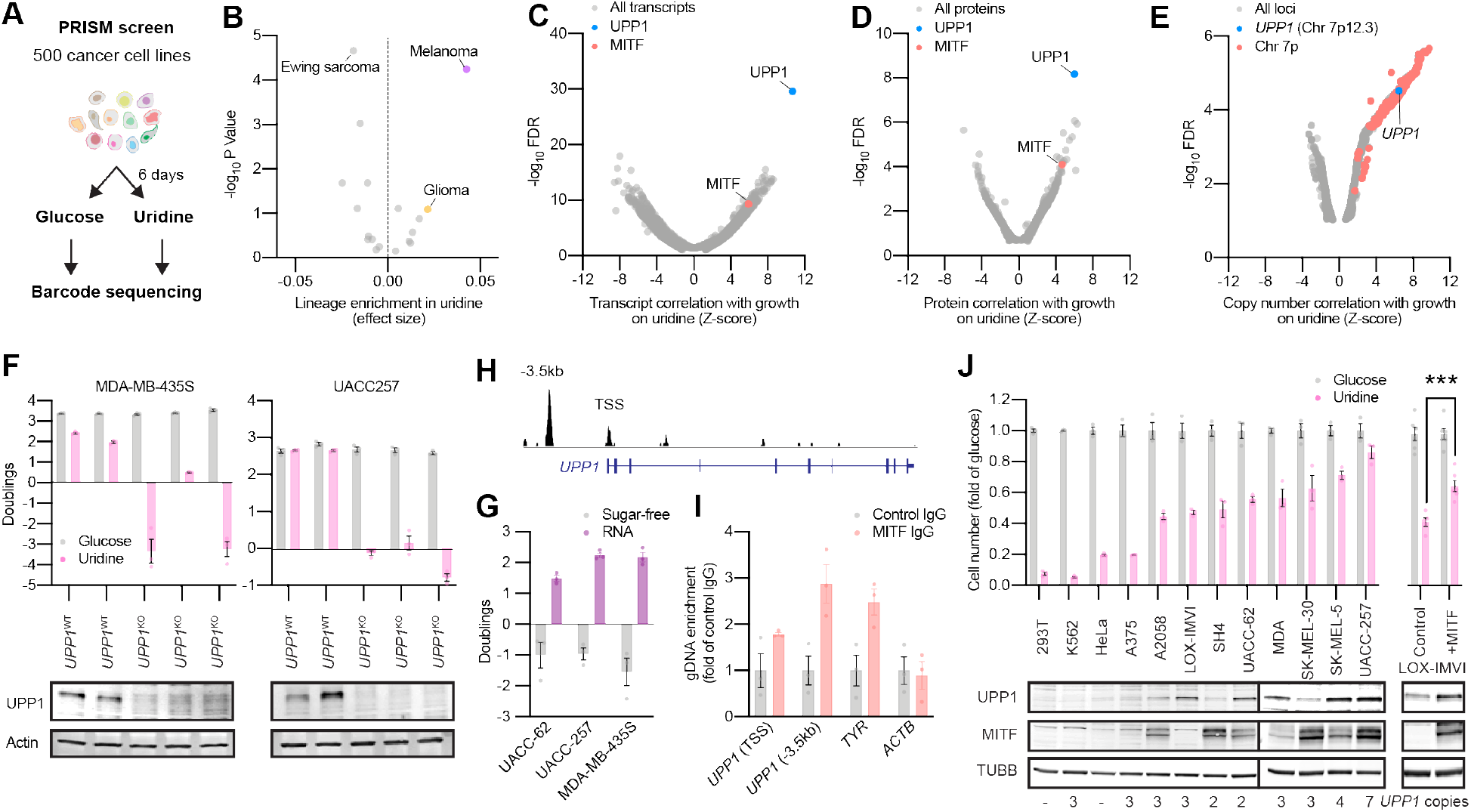
Endogenous *UPP1* Expression Supports Growth of Cancer Cells on Uridine. (A) Schematic representation of the PRISM screen with 482 adherent cancer cells lines grown for 6 days in sugar-free media complemented with 10mM of either glucose or uridine. (B) Lineage analysis (*n*=22 lineages) highlighting increased growth on uridine as compared to glucose after 6 days. (C) Correlation between the ability to grow on uridine and expression of transcript (*n*=8,123) and (D) proteins (*n*=3,216) across the 482 cancer cell lines. (E) Correlation between gene copy number (*n*=5,950) and growth on uridine across the 482 cell lines, highlighting chromosome 7p. *UPP1* is labeled in accordance with its chromosomal location (Chr7p12.3). (F) Cell growth assay and protein immunoblot of two melanoma cell lines (MDA-MB-435S and UACC257) showing two control (*UPP1*^WT^) and three *UPP1*^KO^ cell clones. Negative doublings indicate cell death. Data are shown as mean ±SEM (*n*=3). (G) Cell growth assay of the indicated melanoma cells in sugar-free DMEM complemented with 0.5mg/mL of purified yeast RNA. (H) MITF occupancy in *UPP1* transcription start site (TSS) and promoter (a region 3.5kb away from the TSS), as determined by ChIPSeq in COLO829 melanoma cells(*6*). (I) ChIP-qPCR validation (*n*=3) of MITF binding in *UPP1* promoter and TSS in MDA-MB-435S melanoma cells. *TYR* is a known transcriptional target of MITF. (J) Cell growth assay and protein immunoblot of a panel of melanoma (n=9) and non-melanoma (n=3) cell lines showing global correlation between MITF, UPP1 and growth on uridine. LOX-IMVI is a melanoma cell line with low endogenous MITF expression. Data are shown as mean ±SEM (*n*=3). Growth and qPCR data are shown as mean ±SEM with ***p<10^−3^ and t-test relative to the indicated control.

We next investigated the factors that promote *UPP1* expression and growth on uridine by analyzing our correlation data and prioritizing transcription factors, which highlighted *MITF* (*Microphthalmia associated Transcription Factor*) as a strong candidate, both at the protein and the RNA level. We found binding of this transcription factor in the transcription start site (TSS) and the promoter (−3.5kb from the TSS) of *UPP1* in a large-scale chromatin immunoprecipitation study (*6*), which we could experimentally validate (Fig. 2H-I). Supporting a transcriptional role for MITF in *UPP1* expression, we observed a decrease in UPP1 transcripts in melanoma cells treated with *MITF* siRNA (Fig. S2C), and accordingly MITF, UPP1 and uridine growth correlated with varying MITF levels across a panel of 9 melanoma cell lines and 3 non-melanoma cell lines (Fig. 2J). Furthermore, the abundance of UPP1, as well as the ability to grow on uridine, were further enhanced when we over-expressed MITF in LOX-IMVI, a melanoma cell line with low endogenous MITF expression (Fig. 2J). We conclude that the endogenous expression of *UPP1* is necessary and sufficient to support the growth of cancer cells on uridine and RNA, and that MITF drives its expression in the melanoma lineage.

We next addressed the full molecular and genetic mechanism supporting the growth of UPP1-expressing cells only on uridine in the absence of glucose. We performed a genome-wide CRISPR/Cas9 depletion screen using *UPP1*-FLAG expressing K562 cells grown on glucose or uridine, and performed sgRNA sequencing and gene ontology analysis (Fig. 3A). Many gene sets were found to be essential in either glucose or uridine conditions (Fig. S3, Table S3). However, a comparative analysis identified three major classes of genes that were differentially essential in uridine: [1] As expected, all three enzymes involved in *de novo* pyrimidine synthesis (*CAD*, *DHODH*, *UMPS*) were essential in glucose but dispensable in uridine. Expression of these proteins was reduced in uridine as determined by global proteomics (Fig. 3D, Table S4), consistent with dispensability, while small molecule inhibition of DHODH activity was highly toxic in glucose but not in uridine (Fig. 3E); [2] Genes central to the non-oxPPP (*TKT*, *RPE, PGM2*, *TALDO1*) showed high essentiality in uridine (Fig. 3B-C). In particular, the phosphoglucomutase *PGM2*, which converts ribose-1-P to ribose-5-P and connects the *UPP1/2* reaction to the PPP was highly essential in uridine, but almost fully dispensable in glucose. In contrast, genes of the oxidative branch of the PPP (*G6PD*, *6PGL*, *6PGDH*) did not score differentially between glucose and uridine (Table S3). While we did not observe significant difference in PPP protein abundance in uridine (Fig. 3D), we noticed that uridine grown cells were particularly sensitive to TKT inhibition (Fig. 3E), or to the genetic ablation of *PGM2*, *TKT* or *RPE* (Fig. 3F); [3] As expected from their essentiality in limiting glucose conditions (*7, 8*), genes encoding the mitochondrial respiratory chain were generally more essential in uridine, perhaps due to the high demand in energy in the absence of glucose (Fig. S3), and although to a lesser extent compared to the non-oxPPP.

**Figure 3:**
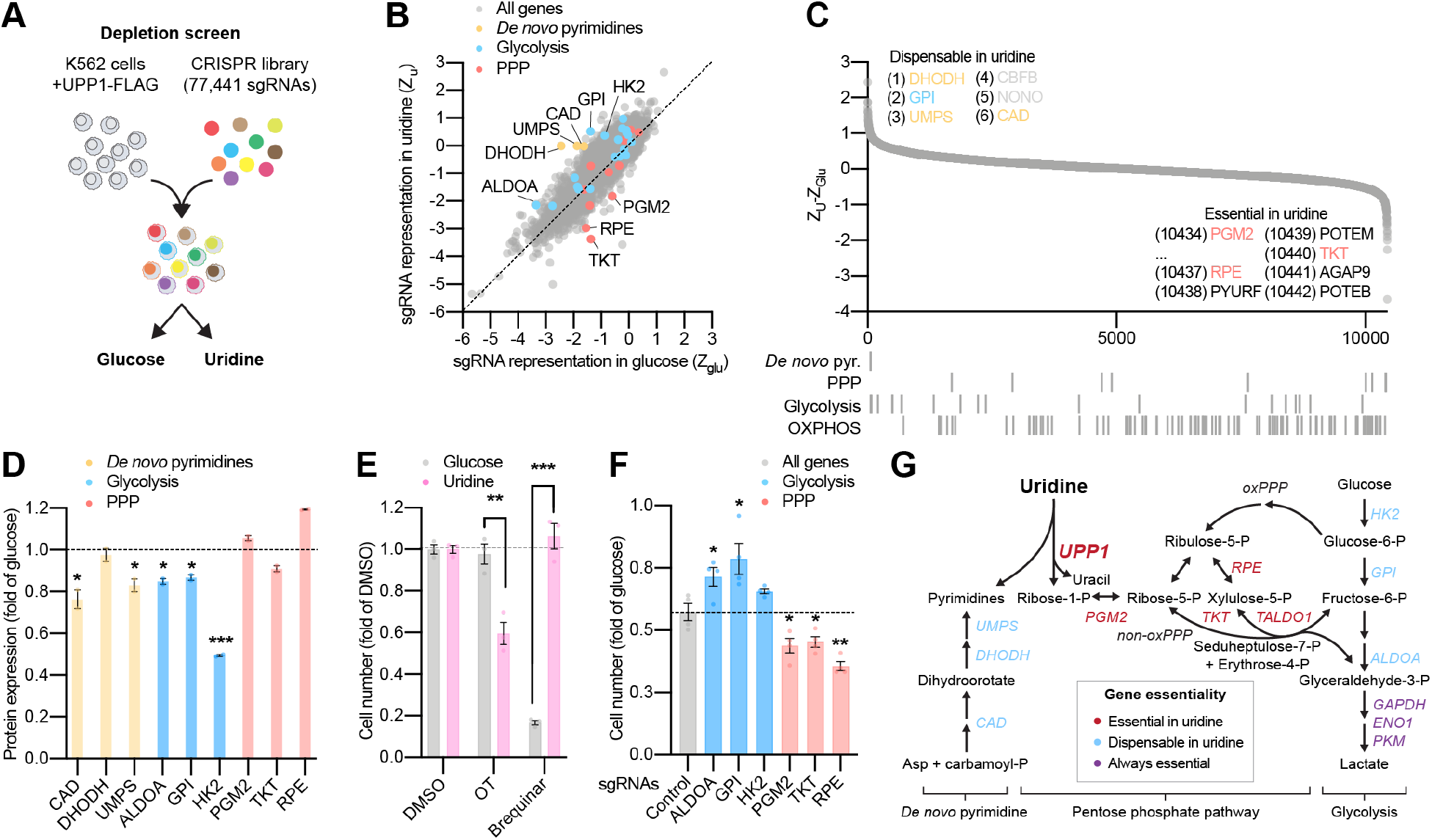
Differential Gene Essentiality in Glucose *vs* Uridine Identifies Key Roles for the PPP and Energy Metabolism Pathways. (A) Schematic representation of the genome-wide CRISPR/Cas9 depletion screen comparing the proliferation of cells in sugar-free media containing 10mM of glucose or 10mM of uridine after 21 days (*n*=2), in the presence of glutamine but in the absence of supplemental pyruvate and uridine. (B) Gene-level analysis of a genome-wide CRISPR/Cas9 screen in glucose *vs* uridine reported as *Z* scores relative to non-cutting controls in glucose (*Z*_Glu_) and uridine (*Z*_U_), or (C) as a ranked list based on (*Z*_U_-*Z*_Gal_) scores. Each dot represents an expressed gene (*n*=10,442), with (n=2) replicates. (D) Abundance of selected proteins from *de novo* pyrimidine synthesis, upper glycolysis and the pentose phosphate pathway (PPP) in cells grown for 15 days in media containing 10mM of either glucose or uridine, as determined by quantitative proteomics (*n*=2). (E) Differential sensitivity to small molecule inhibitors of the PPP (OT: oxythiamine) or *de novo* pyrimidine synthesis (brequinar) in glucose *vs* uridine, reported as fold of DMSO. (F) Differential sensitivity of cells treated with the indicated sgRNAs corresponding to enzymes of upper glycolysis or the PPP in glucose *vs* uridine, expressed as fold of glucose and compared to control sgRNAs. (G) Schematic representation of *de novo* pyrimidine synthesis, the PPP, glycolysis and the results from the differential analysis of gene essentiality in glucose *vs* uridine. Genes highlighted in red and blue are respectively genes essential or dispensable in uridine as compared to glucose. Genes highlighted in gray are essential in both uridine and glucose. Growth and lactate assays are shown as mean ±SEM with *p<0.05, **p<10^−2^ and ***p<10^−3^ t-test relative to the indicated control, with *n*=3-4.

Central enzymes of glycolysis (*GAPDH*, *ENO1*, *PKM*) were essential both in glucose and in uridine (Fig. S3A) indicating that a functional glycolytic pathway is required for the growth of cells in uridine. However, our comparative analysis (Fig. 3B-C) revealed that several upper glycolytic enzymes (*HK2*, *GPI*, *ALDOA*) were dispensable in uridine, but essential in glucose (Fig. 3B-F). Not all steps of upper glycolysis scored in either condition, potentially due to the multiple genes with overlapping functions encoding glycolytic enzymes, a common limitation in single gene-targeting screens. Nevertheless, dispensable genes in uridine all catalyzed steps upstream of fructose-6-P (F6P) and/or glyceraldehyde-3-P (G3P), which connect the non-oxPPP to glycolysis, pointing to a key role for these two metabolites in supporting proliferation on uridine. Together, our data is compatible with a model in which uridine feeds into glycolysis via the non-oxPPP, by a mechanism that appears to bypass upper glycolysis (Fig. 3G).

To test this model, we investigated the bioenergetics of uridine growth. Cultured cells typically draw equivalent parts of their ATP from glucose and glutamine (*9*), and most cells can tolerate a loss of glucose and compensate by oxidizing glutamine (*7*). We thus measured how respiration (oxygen consumption rate, OCR) and glycolysis (extracellular acidification rate, ECAR) were affected in uridine. We found that, independent of *UPP1* expression, mitochondrial respiration was maintained in glucose, uridine, galactose or sugar-free media (Fig. 4A), suggesting maintenance of respiration by glutamine. However, we found a strong difference in ECAR: whereas in control cells only glucose supported high ECAR (Fig. 4B), we observe that uridine can support high ECAR in a *UPP1*-dependent manner (Fig. 4B right). Interestingly, unlike with glucose, we did not observe regulation of glycolytic flux by inhibition of the respiratory chain in uridine (Fig. 4B). We further tested for the presence of lactate in the culture media of these cells and found significant levels in *UPP1*-expressing cells in uridine (Fig. 4C). Thus, uridine and *UPP1* can support glycolysis in the absence of glucose.

**Figure 4:**
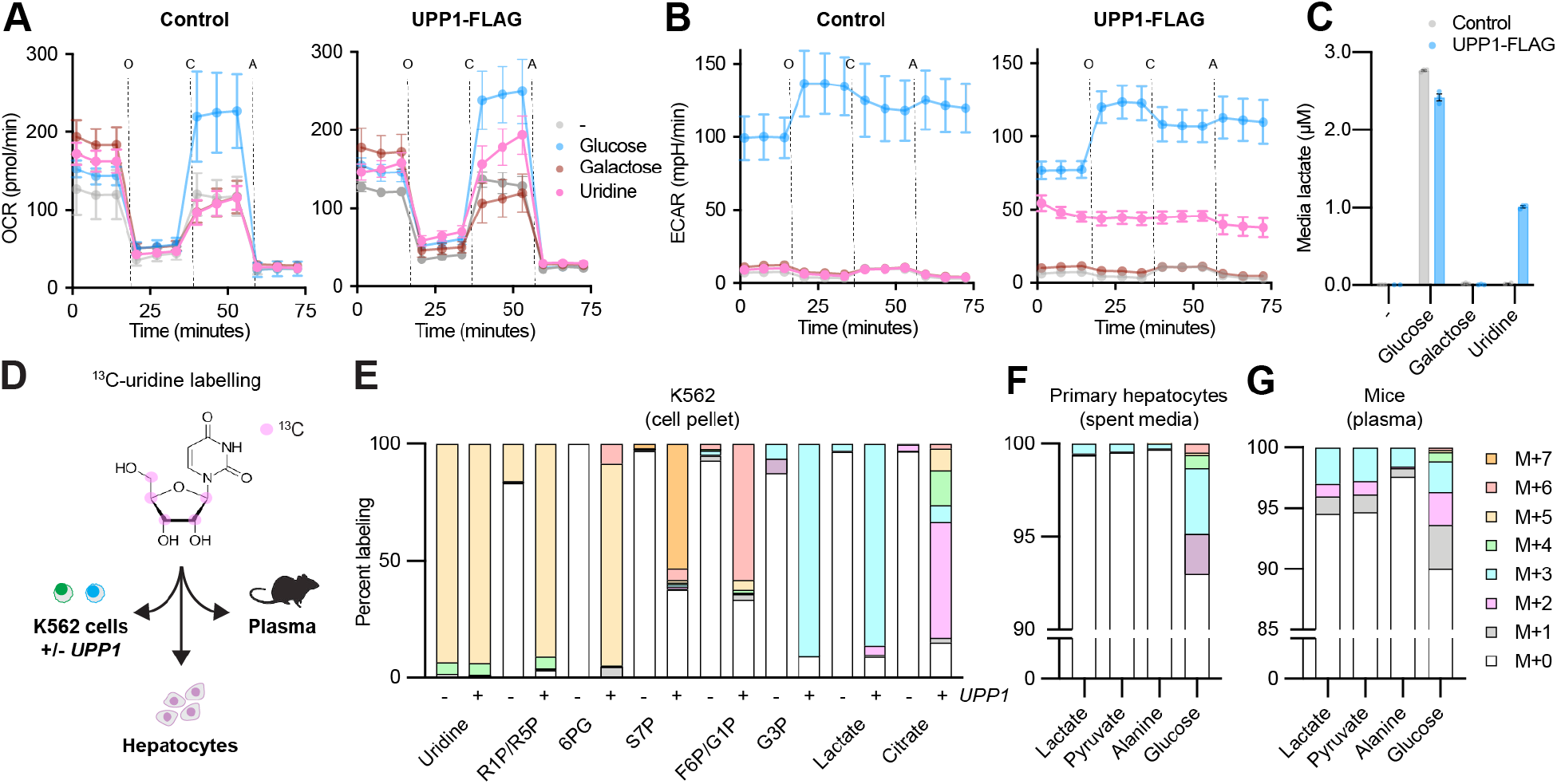
Uridinolysis Fuels Glycolysis, the TCA Cycle and Gluconeogenesis in Cells and Animals. (A) Representative oxygen consumption rate (OCR) and (B) extracellular acidification rate (ECAR) in K562 cells expressing a control gene or *UPP1* and grown in sugar-free media or in media supplemented with 10mM of glucose, galactose or uridine, all in the presence of glutamine. Data are shown as mean ±SD with *n*>6 replicate wells. (C) Media lactate determination in the same conditions after three hours (*n*=3). Data are shown as mean ±SEM. (D) Schematic representation of our labelling strategy with ^13^C-uridine ([1′,2′,3′,4′,5′-13C_5_]uridine). Labeled carbon atoms in the ribose of uridine are indicated in magenta. (E) ^13^C-uridine tracer analysis reporting representative intracellular metabolites from the PPP, glycolysis and the TCA cycle in K562 cells expressing *GFP* or *UPP1*-FLAG after 5h (*n*=4). R1P/R5P: ribose-1/5-P; 6PG: 6-phosphogluconate; S7P: sedoheptulose-7-P; F6P/G1P: fructose-6-P/glucose-1-P. G3P: glyceraldehyde-3-P. Data are shown as percent labelled fraction after correction for natural isotope abundance (*n*=3). (F) ^13^C-uridine tracer analysis reporting representative secreted metabolites related to glycolysis/gluconeogenesis in primary mouse hepatocytes after 5h in sugar-free DMEM media, in the absence of supplemental pyruvate. Data are shown as percent labelled fraction after normalization for natural isotope abundance (*n*=3-4). (G) ^13^C-uridine tracer analysis reporting representative circulating metabolites related to glycolysis/gluconeogenesis in mouse plasma 30min after intraperitoneal (IP) injection with PBS or 0.4g/kg ^13^C-uridine. Data are shown as percent labelled fraction after correction for natural isotope abundance (*n*=4).

The crucial role of *PGM2* for the growth of cells on uridine prompted us to hypothesize that the ribose moiety of uridine can enter glycolysis. To directly test this hypothesis, we designed a tracer experiment using isotopically labeled uridine with five ribose carbons labeled (^13^C_5_-uridine) and liquid chromatography-mass spectrometry (LC-MS) (Fig. 4D). UPP1-expressing K562 cells avidly incorporated ^13^C_5_-uridine, as seen by the presence of ^13^C in all the intracellular intermediates of the PPP and glycolysis analyzed, including ribose-phosphate, upper and lower glycolytic intermediates and lactate, while control cells showed very little label incorporation (Fig. 4E). TCA cycle intermediates, among them citrate, were also partially labelled (mostly M+2), indicating origin of carbon from glycolysis via pyruvate. We next repeated our LC-MS experiments using non-labelled uridine and compared steady-state abundance of central carbon metabolism with other sugars. As expected from the enzymatic activity of UPP1, we found a drastic accumulation of ribose-P in uridine, but also of PPP and glycolytic intermediates, confirming communication between the UPP1 reaction, the PPP and glycolysis (Fig. S4). Together, our cell culture experiments indicate that the phosphorylytic cleavage of uridine provides ribose for the PPP, and that the non-oxPPP and the glycolytic pathway communicate via F6P and G3P to replenish glycolysis, thus bypassing the requirement of glucose in supporting lower glycolysis, biosynthesis and energy production in sugar-free media.

We next tested whether ^13^C_5_-uridine could be metabolized in primary cells and in animals. In mice, uridine is primarily catabolized by UPP1/2 the liver (*10*), maintaining circulating uridine levels to 5-20μM (*10*). Fasting promotes expression of these genes (*11*) as well as gluconeogenesis and we tested how primary hepatocytes grown in sugar-free conditions would process ^13^C-labelled uridine. We found that these cells were able to secrete a significant fraction (∼7%) of ^13^C-labelled glucose as well as a smaller fraction of ^13^C-labeled alanine, ^13^C pyruvate and ^13^C lactate (Fig. 4F). Labelling in glucose was mostly M+3, indicating entry into gluconeogenesis via a 3-carbon molecule, such as glyceraldehyde-3-P, and a large participation from unlabeled substrates, such as amino-acids which are ordinarily the substrates for hepatocyte gluconeogenesis *ex-vivo (12)*. Finally, we sought to determine whether uridine can serve as a glycolytic and/or gluconeogenic substrate *in vivo*. To this end, we injected fasted mice intraperitoneally with ^13^C_5_-uridine and measured incorporation in circulating metabolites after 30min. Similar to cell culture, we found ^13^C in alanine, pyruvate, lactate and up to ∼11% in plasma glucose (Fig. 4G), indicating rapid uridinolysis in these animals. Together, we conclude that uridine can serve as a substrate for glycolysis and gluconeogenesis *in viv*o.

It has long been known that cells with mitochondrial deficiencies are dependent on uridine to support pyrimidine synthesis given the dependence of *de novo* pyrimidine synthesis on DHODH which is coupled the electron transport chain (*1*). However, it is less appreciated that uridine supplementation can support growth of cells with intact electron transport chain in the absence of glucose (*13*). Here, we show that, in addition to nucleotide synthesis, uridine can serve as a substrate for glycolytic ATP and gluconeogenesis. Mechanistically, we show that uridinolysis is initiated with the phosphorylytic cleavage of uridine by UPP1/2, shuttling of its ribose moiety through the non-oxPPP and glycolysis, hence supporting not only nucleotide metabolism but also energy production or gluconeogenesis in the absence of glucose (Fig. 3G).

By comparing uridine to other nucleosides and using similar tracer experiments to ours, Kennell *et al.* (*13*) observed incorporation of uridine-derived carbons in most cellular factions in mammalian cell culture and in chicken embryos. However, they did not detect pyruvate and lactate in uridine, and concluded that uridine do not participate to glycolysis, but rather is required for nucleotide synthesis (*13, 14*). Our observations agree with prior observations that cells can proliferate in sugar-free media if uridine is provided, and uridine is crucial for nucleotide synthesis – but differ mechanistically on the role of glycolysis in this condition, as we were able to identify significant amount of labelling in glycolytic intermediates, lactate production and high ECAR, all consistent with glycolytic ATP production in uridine.

Our work highlighting that uridine breakdown can represent an unregulated, alternative input into glycolysis might have clinical implications. Previous studies have shown that *in vivo* a uridine-rich diet leads to glycogen accumulation, gluconeogenesis, fatty liver and pre-diabetes in mice (*15, 16*), while acute uridine injection sharply increased glycemic index (*17*). Uridine is the most abundant, circulating nucleoside (*18*), and foods such as milk and beer are particularly rich in uridine (*19, 20*). Transport of glucose into cells is tightly regulated, and we propose that “uridine bypass” of this controlled step leads to uncontrolled glycolysis and gluconeogenesis from a uridine-rich diet may contribute to human conditions such as diabetes. It is noteworthy that, unlike with glucose, inhibition of the respiratory chain in uridine condition did not upregulate glycolytic flux (Fig. 4B). This is likely because the regulation of glycolysis in response to AMPK activation occurs largely at the level of upper glycolysis, perhaps at the level of glucose transport (*21*), which we show is bypassed by uridine, since the non-oxPPP and glycolysis are connected by F6P and G3P (Fig. 4H). The ability of uridine to bypass upper glycolysis, as shown in our secondary CRISPR screen (Fig. 3), may be beneficial in certain cases. For example, disorders of upper glycolysis, such as low glucose transport in GLUT1 deficiency syndrome (*22*), may benefit from high *UPP1/2* expression and uridinolysis, and accordingly uridine protects cortical neurons from glucose-deprivation-induced cell death (*23*), and our work suggests that it is likely because it supports glycolytic ATP production and biosynthesis in these cells. We propose that supplementation with dietary uridine may prove to be beneficial in GLUT deficiencies and other pathologies characterized by low glucose availability.

Our profiling effort on cancer cells showed that some cancer cell lines, in particular those of the melanoma and glioma lineages, express high levels of *UPP1* and present a high uridine utilization potential. It is possible that uridine may serve as an alternative energy source in solid tumors as what has been proposed for thymidine (*24*). In this context, uridine may generate glycolytic ATP in glucose-limited conditions, such as in the tumor micro-environment. Supporting this, *UPP1*-depleted xenografts present strong growth defects *in vivo* in mice, while they grow at a rate comparable to wild-type cells in glucose-rich, uridine-free cell culture conditions (*25*). Nucleotide metabolism represents an important target in cancer therapy, with drugs such as brequinar targeting pyrimidine synthesis along to pyrimidine analogs such as 5-fluorouracil (5-FU) being at the front line of treatment for certain cancers. Our findings suggest that the catabolism of uridine and its shunting into glycolysis plays an important role in this context. For example, the metabolism of 5-FU into a potent pyrimidine analog requires the addition of R1P to 5-FU in an anabolic reaction facilitated by *UPP1/2* (*26*). Because we showed that, in low glucose conditions, the UPP1/2 reaction goes in the catabolic direction and ribose-1-P is generated, we predict that high circulating glucose will improve the efficacy of 5-FU in its anti-cancer activity.

We have also found that that RNA in the media can replace glucose to promote cellular proliferation (Fig. 1F, 2G). RNA is a highly abundant molecule, representing from 4% of the dry weight of a mammalian cell to 20% of a bacterium (*27*). Recycling of ribosomes through ribophagy for example plays an important role in supporting viability during starvation (*28*), and cells of our immune system presumably ingest large quantities of RNA during phagocytosis (*27*). Whereas the salvage of RNA to provide building blocks during starvation has long been appreciated, to our knowledge its contribution to energy metabolism has not been considered in the past, despite the fact that certain unicellular organisms can grow on minimum media with RNA as their sole carbon source (*29*). We propose that, similar to glycogen and starch, RNA itself may constitute as large stock of energy under the form of a polymer, and that it may be used for energy storage and to support cellular function during starvation.

## Acknowledgements

The authors would like to thank Olga Goldberger, Danny Rosenberg, and Melissa Ronan for technical assistance. This work was supported by NIH grants R35GM122455 (V.K.M.), F32GM133047 (O.S.S.), DK115881 (R.P.G), R01AR043369-24 (D.E.F.), P01CA163222-07 (D.E.F.), K99/R00 GM124296 (H.S.), an EMBO long-term ALTF 554-2015 fellowship (A.A.J.), an SNF Advanced Postdoc.Mobility P300PA_171514 fellowship (A.A.J.), and a grant from the Dr. Miriam and Sheldon Adelson Medical Research Foundation (to D.E.F.). V.K.M. is an Investigator of the Howard Hughes Medical Institute.

## Methods

### Cell Lines

K562 (CCL-243), 293T (CRL-3216), HeLa (CCL-2), A375 (CRL-1619), A2058 (CRL-11147), SH4 (CRL-7724), MDA-MB-435S (HTB-129), SK-MEL-5 (HTB-70) and SK-MEL-30 (HTB-63) were obtained from ATCC. UACC-62, UACC-257 and LOX-IMVI were obtained from the Frederick Cancer Division of Cancer Treatment and Diagnosis (DCTD) Tumor Cell Line Repository. All cell lines were re-authenticated by STR profiling at ATCC prior submission of the manuscript and compared to ATCC and Cellosaurus (ExPASy) STR profiles.

## METHOD DETAILS

### Cell Culture and Cell Growth Assays

Cell lines stocks were routinely maintained in DMEM media (HeLa, 293T, K562, A375, A2058, SK-MEL-5, MDA-MB-435S) containing 1mM sodium pyruvate (ThermoFisher Scientific) with 25 mM glucose, 10% fetal bovine serum (FBS, ThermoFisher Scientific), 50 μg/mL uridine (Sigma), and 100 U/mL penicillin/streptomycin (ThermoFisher Scientific) ; or in RPMI (SH4, UACC-62, UACC-257, SK-MEL-30, LOX-IMVI) containing 10% fetal bovine serum (FBS, ThermoFisher Scientific) and 100 U/mL penicillin/streptomycin (ThermoFisher Scientific), under 5% CO_2_ at 37°C. For growth experiments, an equal number of cells was counted, washed in PBS and resuspend in no glucose DMEM (ThermoFisher Scientific, or no glucose RPMI (Teknova) complemented with 10% dialyzed fetal bovine serum (FBS, ThermoFisher Scientific), 100 U/mL penicillin/streptomycin (ThermoFisher Scientific) and 5-10mM of either glucose, galactose or uridine (all from Sigma) dissolved in water, or with an equal volume of water alone. For RNA and other nucleoside complementation assays, purified RNA from *Torula* yeast (Sigma) or the selected nucleosides were weighted and directly resuspend in DMEM before sterile filtration. In all cases, cells were counted with a ViCell Counter (Beckman) after 3-4 days of growth and only live cells were considered.

### ORFeome Screen

For ORFeome screening, K562 cells were infected with a lentiviral-carried ORFeome v8.1 library (Genome Perturbation Platform, Broad Institute) containing 17,255 ORFs mapping to 12,826 genes (*2*), in duplicate. Cells were infected with multiplicity of infection (MOI) of 0.3 and at 500 cells/sgRNA in the presence of polybrene (Millipore). After 72h, cells were transferred to culture media containing 2 mg/mL puromycin (ThermoFisher Scientific) and incubated for an additional 48h. On day of the screen, cells were plated in screening media containing no glucose DMEM supplemented with 10% FBS, 1mM sodium pyruvate (ThermoFisher Scientific), 50 μg/mL uridine (Sigma), and 100 U/mL penicillin/streptomycin (ThermoFisher Scientific) and 25mM of either glucose or galactose (Sigma) at a concentration of 10^5^ cells/mL and with 500 cells/ORFs. Cells were passaged every 3 days and 500 cells/ORFs were harvested after 0, 9 and 21 days of growth. Total genomic DNA was isolated from cells using a NucleoSpin Blood kit (Clontech) using the manufacturer’s recommendations. Barcode sequencing, mapping and read-counts were performed by the Genome Perturbation Platform (Broad Institute). Log_2_ normalized read counts were used for screen analysis, and P values were calculated using a T test.

### Stable Gene Over-Expression

cDNAs corresponding to *GFP*, *UPP1*-FLAG and *UPP2-*FLAG were cloned in pWPI/Neo (Addgene). Lentiviruses were produced according to Addgene’s protocol. 24h post-infection, cells were selected with 0.5 mg/mL Geneticin (ThermoFisher Scientific) for 48h.

### Polyacrylamide Gel Electrophoresis and Immunoblotting

Cells were harvested, washed in PBS and lysed for 5min on ice in RIPA buffer (25mM Tris pH 7.5, 150mM NaCl, 0.1% SDS, 0.1% sodium deoxycholate, 1% NP40 analog, 1x protease (Cell Signaling) and 1:500 Universal Nuclease (ThermoFisher Scientific). Protein concentration was determined from total cell lysates using DC protein assay (Biorad). Gel electrophoresis was done on Novex Tris-Glycine gels (ThermoFisher Scientific) before transfer using the Trans-Blot Turbo blotting system and nitrocellulose membranes (Biorad). All immunoblotting was performed in Intercept Protein blocking buffer (Licor). Washes were done in TBS + 0.1% Tween-20 (Sigma). Specific primary antibodies were diluted 1:100-1:5000 in blocking buffer. Fluorescent-coupled secondary antibodies were diluted 1:10,000 in blocking buffer. Membranes were imagined with an Odyssey CLx analyzer (Licor) or by chemiluminescence. The following antibodies were used: FLAG M2 (Sigma, F1804), Actin (Abcam, ab8227), UPP1 (Sigma, SAB1402388) and MITF (Sigma, HPA003259).

### PRISM Screen

A 6-well plate containing a mixture of 482 barcoded adherent cancer cell lines (PR500) (*3*) grown on RPMI (Life Technologies, 11835055) containing 10% FBS was prepared by the PRISM lab (Broad Institute) seeded at a density of 200 cells per cell line. On day 0, the culture media was replaced for no glucose RPMI media (Life Technologies, 11879020) containing 10% dialyzed FBS and 100 U/mL penicillin/streptomycin and supplemented with 10mM of either glucose or uridine (*n*=3 replicate wells each). The media was replaced with fresh media on days 3 and 5. On day 6, all wells reached confluency and cells were lysed. Lysates were denatured (95°C) and total DNA from all replicate wells was PCR-amplified using KAPA polymerase and primers containing Illumina flow-cell binding sequences. PCR products were confirmed to show single-band amplification using gel electrophoresis, pooled, purified using the Xymo Select-a-Size DNA Clean & Concentration kit, quantitated using a Qubit 3 Fluorometer, and then sequenced via HiSeq (50 cycles, single read, library concentration 10pM with 25% PhiX spike-in) as previously described (*30*). Barcode abundance was determined from sequencing, and unexpectedly low counts (e.g. from sequencing noise) were filtered out from individual replicates so as not to unintentionally depress cell line counts in the collapsed data. Replicates were then mean-collapsed, and log-fold change and growth rate metrics were calculated as follows:

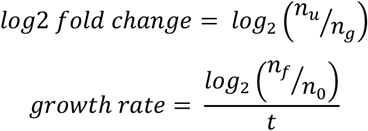

where n_u_ and n_g_ are counts from the uridine and glucose supplemented conditions, respectively, n_0_ and n_f_ are counts from the initial and final timepoints, respectively, and t is the assay length in days. Data analysis and correlation analysis were performed by the PRISM lab following a published workflow (*3*).

### Chromatin-Immunoprecipitation

MDA-MB-435S cells were washed once with PBS and fixed with 1% formaldehyde in PBS for 15 min at room temperature. Fixation was stopped by adding glycine (final concentration of 0.2 M) for 5 min at room temperature. Cells were harvested by scraping with ice cold PBS. Cell pellets were resuspended in SDS lysis buffer (50 mM Tris-HCl, pH8.1, 10 mM EDTA, 1% SDS, protease inhibitor (Pierce™ Protease Inhibitor, EDTA-Free (Thermo Fisher Scientific))), incubated for 10 min at 4 °C, and sonicated to make DNA fragments (around 500 base pairs) with Qsonica Q800R2 system (Qsonica). Samples were centrifuged to remove debris and diluted 10-folds in IP dilution buffer (16.7 mM Tris-HCl, pH 8.1, 1.2 mM EDTA, 0.01% SDS, 1.1% Triton-X100, 167 mM NaCl, protease inhibitor).

Chromatin (∼50 μg) was pre-cleared with normal rabbit IgG (EMD Millipore) and protein A/G beads (Protein A/G UltraLink® Resin (Thermo Fisher Scientific)) in low salt buffer (20 mM Tris-HCl, pH8.1, 2 mM EDTA, 0.1% SDS, 150 mM NaCl, protease inhibitor) containing 0.25 mg/ml salmon sperm DNA and 0.25 mg/ml BSA for 2 h at 4 °C. Pre-cleared chromatin was incubated with 5 μl of anti-MITF antibody (D5G7V (Cell Signaling Technology)) or 5 μg of normal rabbit IgG overnight at 4 °C. Samples were incubated with protein A/G beads for another 2 h at 4 °C. Immune complexes were washed sequentially twice with low salt buffer, twice with high salt buffer (20 mM Tris-HCl, pH8.1, 2 mM EDTA, 0.1% SDS, 500 mM NaCl, protease inhibitor), LiCl buffer (250 mM LiCl, 1% NP40, 1% sodium deoxycholate, 1 mM EDTA, 10 mM Tris-HCl, pH 8.1, protease inhibitor), and twice with TE. After washes, immune complexes were eluted from beads twice with elution buffer (1% SDS, 10 mM DTT, 0.1M NaHCO_3_) for 15 min at room temperature. Samples were decross-linked by overnight incubation at 65 °C and treated with proteinase K (Qiagen) for 1 h at 56 °C. DNA was purified with QIAquick PCR purification kit (Qiagen).

qPCR using KAPA SYBR® FAST One-Step qRT-PCR Kit Universal (KAPA Biosystems) was performed to check MITF enrichments using the following primers: *UPP1*-TSS (5’-TGACCTTGGGTTAGTCCTAGA-3’) and (5’-AGCAGCCAGTTCTGTTACTC-3’) ; *UPP1*—3.5kb (5’-AGCAACCTGGGAAAGTGATG-3’) and (5’-CGCCAACTCTCACTCATCATATAG-3’) ; *TYR* promoter (5’-GTGGGATACGAGCCAATTCGAAAG-3’) and (5’-TCCCACCTCCAGCATCAAACACTT-3’) ; *ACTB* gene body (5’-CATCCTCACCCTGAAGTACCC-3’) and (5’-TAGAAGGTGTGGTGCCAGATT-3’)

### Gene-Specific CRISPR-Cas9 Single Cell Clone Knockouts

An sgRNAs targeting *UPP1* (TTGGATTTAAAAGTCTGACG) was ordered as complementary oligonucleotides (Integrated DNA Technologies) and cloned in pLentiCRISPRv2. Purified DNA was co-transfected with a GFP-expressing plasmid in the cell lines of interest using Lipofectamine 2000 (ThermoFisher Scientific). After 48h, cells were sorted using an MoFlo Astrios EQ Cell sorter and individual cells were seeded in a 96-well plate containing routine culture media for clone isolation. *UPP1*-depletion in single cell clones was assessed by protein immunoblotting using antibodies to UPP1.

### siRNA Treatment

UACC-257 and MDA-MB-435S were transfected with a non-targeting siRNA (N-001206-14-05) or an siRNA targeting MITF (M-008674-0005) (Dharmacon) using Lipofectamine RNAiMAX according to the manufacturer’s instruction. Cells were analyzed 72h post-transfection and significant *MITF* knock-down was confirmed by qPCR.

### RNA-Extraction, Reverse Transcription and qPCR

qPCR was performed using the TaqMan assays (ThermoFisher Scientific). RNA was extracted from total cells with a RNeasy kit (QIAGEN) and DNase-I digested before murine leukemia virus (MLV) reverse transcription using random primers (Promega) and a CFx96 quantitative PCR machine (Biorad) using Taqman assays. All data were normalized to *TBP* using ΔΔCt method. Taqman assays were Hs01066247_m1 (*UPP1*), Hs01117294_m1 (*MITF*) and Hs00427620_m1 (*TBP*).

### Genome-Wide CRISPR/Cas9 Screening

A secondary genome-wide CRISPR/Cas9 screening was performed using K562 cells expressing *UPP1*-FLAG and a lentiviral-carried Brunello library (Genome Perturbation Platform, Broad Institute) containing 76,441 sgRNA (*31*), in duplicate. Cells were infected with multiplicity of infection (MOI) of 0.3 and at 500 cells/sgRNA in the presence of polybrene (Millipore). After 24h, cells were transferred to culture media containing 2 mg/mL puromycin (ThermoFisher Scientific) and incubated for an additional 48h. On day 7, the cells were plated in no glucose DMEM containing 10% FBS and 100 U/mL penicillin/streptomycin and supplemented with 10mM of either glucose or uridine at a concentration of 10^5^ cells/mL and with 1000 cells/sgRNA. Cells were passaged every 3 days for 2 weeks and on day 21 1000 cells/sgRNA were harvested. DNA isolation was performed as for the ORFeome screen.

CRISPR screen analysis was performed using a normalized Z-score approach where raw sgRNA read counts were normalized to reads per million and then log_2_ transformed using the following formula: Log_2_(reads from an individual sgRNA / total reads in the sample10^6^ + 1) (*32*). Log_2_ fold-change of each sgRNA was determined relative to the pre-swap control. For each gene in each replicate, the mean log_2_ fold-change in the abundance of all 4 sgRNAs was calculated. Genes with low expression (log_2_ FPKM<0) according to publicly available K562 RNA-seq dataset (sample GSM854403 in GEO seriesGSE34740) were removed. Log_2_ fold-changes were averaged by taking the mean across replicates. For each treatment, a null distribution was defined by the 3,726 genes with lowest expression. To score each gene within each treatment, its mean log_2_ fold-change across replicates was Z-score transformed, using the statistics of the null distribution defined as above.

### Quantitative Proteomics

Samples were generated at the same time as our analysis of the proteome remodeling in galactose (*33*) and are available at the PRIDE consortium. K562 stably expressing *UPP1*-FLAG cells were grown in duplicate plates in no glucose DMEM media containing 10mM of either glucose or uridine for 2 weeks. Quantitative proteomics was performed at the Thermofisher for Multiplexed Proteomics (Harvard). In short, cells were harvested and total protein quantification was performed using micro-BCA assay (Pierce). Samples were reduced with DTT and alkylated with iodoacetamide before protein precipitation in methanol/chloroform. Pellets were resuspended in 200 mM EPPS, pH 8.0 and a digestion was performed sequentially using LysC (1:50) and Trypsin (1:100) based on protease to protein ratio. ∼50μg peptide per sample was labeled with TMTpro 16 reagents. A small aliquot of each sample was then combined and analyzed by LC-MS3 to verify labeling efficiency and mixing ratios. Samples were combined, desalted, and dried by speedvac. 14 fractions from the total proteome HPRP set were analysed on an Orbitrap Eclipse mass spectrometer using a 180-minute method MS3 method with real-time search. Peptides were detected (MS1) and quantified (MS3) in the Orbitrap. Peptides were sequenced (MS2) in the ion trap. MS2 spectra were searched using the COMET algorithm against a custom protein database containing only one protein per gene (referenced as the canonical isoform). Peptide spectral matches were filtered to a 1% false discovery rate (FDR) using the target-decoy strategy combined with linear discriminant analysis. The proteins from the 14 runs were filtered to a <1% FDR. Proteins were quantified only from peptides with a summed SN threshold of >100. Only unique peptides were considered for downstream analysis. The mass spectrometry proteomics data have been deposited to the ProteomeXchange Consortium via the PRIDE (*34*) partner repository with the dataset identifier PXD021917 and 10.6019/PXD021917”.

### Metabolite Profiling

For steady-state metabolomics for glycolytic and PPP intermediates, an equal number of K562 cells expressing *GFP* or *UPP1*-FLAG were washed in PBS pre-incubated for 24h in no glucose DMEM media supplemented with 10% dialyzed fetal bovine serum (FBS, ThermoFisher Scientific), 100 U/mL penicillin/streptomycin (ThermoFisher Scientific) and 5mM of either glucose, galactose or uridine (all from Sigma) dissolved in water, or with an equal volume of water alone. Cells were then re-counted and 2 × 10^6^ cells were seeded in fresh media of the same formulation and incubated for two additional hours before metabolite extraction Cells were pelleted and immediately extracted with 80% MeOH, lyophilized and resuspended in 60% acetonitrile for intracellular LC-MS analysis.

### ^13^C-Uridine Tracer on Cultured Cells

For tracer analysis on cultured cells, an equal number of K562 cells expressing *GFP* or *UPP1*-FLAG were washed in PBS pre-incubated for 24h in no glucose DMEM media supplemented with 10% dialyzed fetal bovine serum (FBS, ThermoFisher Scientific), 100 U/mL penicillin/streptomycin (ThermoFisher Scientific) and 5mM of either glucose or unlabeled uridine (all from Sigma) dissolved in water, or with an equal volume of water alone. Cells were then re-counted and 7 × 10^6^ cells were seeded in fresh media of the same formulation, with the exception of cells pre-incubated in uridine that were either grown on unlabeled uridine, or on ^13^C-labelled uridine ([1′,2′,3′,4′,5′-13C5]uridine, NUC-034, Cambridge Isotope Laboratories), and incubated for five additional hours before metabolite extraction. Cells were then harvested, the media was removed, and cellular pellets were resuspending in an 9:1 ((75% acetonitrile; 25% MetOH):water)) extraction mixture, spun at 20,000 × g for 10 minutes, and the supernatant was transferred to a glass sample vial for LCMS analysis.

### ^13^C-Uridine Tracer on Primary Hepatocytes

Primary mouse hepatocytes were isolated via collagenase perfusion of liver of male C57BL/6J mice and plated on six-well collagen-coated plates (A1142801, Life Technologeis) at a density of 4 × 10^5^ cells/well in DMEM media (11995-065, Gibco) supplemented with 10% FBS (F2442, Sigma-Aldrich) and 100 U/mL penicillin/streptomycin. On the day of the experiment, the culture media was replaced with no sugar DMEM complemented supplemented with 10% dialyzed fetal bovine serum (FBS, ThermoFisher Scientific), 100 U/mL penicillin/streptomycin (ThermoFisher Scientific) and incubated for 4h for glucose starvation. Cells were then grown in no glucose DMEM media supplemented with 10% dialyzed fetal bovine serum (FBS, ThermoFisher Scientific), 100 U/mL penicillin/streptomycin (ThermoFisher Scientific) and 5mM of either unlabeled uridine or ^13^C-labelled uridine (1′,2′,3′,4′,5′-13C5]uridine dissolved in water, or with an equal volume of water alone, and incubated for an additional 5h before media collection and metabolite extraction. For metabolite extraction, 20 uL of LCMS-grade water and 117 uL of acetonitrile were added to 30 uL of each sample, the mixture was vortexed and left on ice for 10 minutes, and then spun at 21,000 × g for 20 minutes. 100 uL of the supernatant was transferred to a glass sample vial and subjected to LCMS analysis

### ^13^C-Uridine Tracer in Mice

For tracer analysis on cultured cells, male 8-12 week old C57BL/6 mice were fasted overnight and injected intraperitoneally with 200uL of 0.2M ^13^C-labelled uridine diluted in PBS, or with PBS alone. After 30min, blood was collected from mice under isoflurane anesthesia. For plasma metabolite analysis, 117 uL of acetonitrile and 20 uL of LCMS-grade water were added to 30 uL of plasma, the mixture was vortexed and left on ice for 10 minutes. The samples were then spun at 21,000 × g for 20 minutes, and 100 uL of the supernatant was transferred to a glass sample vial for downstream LCMS analysis.

### Intracellular LC-MS Analysis

For labeled and unlabeled LCMS analysis of intracellular metabolites, 5 uL of sample was loaded on a ZIC pHILIC column (Milipore). Buffer A was 20 mM ammonium carbonate, pH 9.6 and Buffer B was acetonitrile. For each run, the total flow rate was 0.15 mL/min and the samples were loaded at 80%B. The gradient was held at 80% B for 0.5 minutes, then ramped to 20% B over the next 20 minutes, held at 20% B for 0.8 minutes, ramped to 80% B over 0.2 minutes, then held at 80% B for 7.5 minutes for re-equilibration. Mass spectra were continuously acquired on a Thermo Q-Exactive Plus run in polarity switching mode with a scan range of 70-1000 *m/z* and a resolving power of 70,000 (@200 *m/z*). Data was analyzed using Tracefinder software, and labeled data was manually corrected for natural isotope abundance.

### Media/Plasma LC-MS Analysis

Media and plasma samples were subjected to the following LC-MS analysis: 10 uL of sample was loaded on a BEH Amide column (Waters). Buffer A was 20 mM ammonium acetate, 0.25% ammonium hydroxide, 5% acetonitrile, pH 9.0, while buffer B was acetonitrile. Samples were loaded on the column and the gradient began at 85% B, 0.22 mL/min, held for 0.5 minutes, then ramped to 35% B over 8.5 minutes, then ramped to 2% B over two minutes, held for one minute, then ramped to 85% B over 1.5 minutes and held for 1.1 minutes. The flow rate was then increased to 0.42 mL/min and held for 3 minutes for re-equilibration. Mass spectra were collected on a Thermo Q-Exactive Plus run in polarity switching mode with a scan range of 70-1000 *m/z* and a resolving power of 70,000 (@200 *m/z*). Data was analyzed using Tracefinder software, and labeled data was manually corrected for natural isotope abundance.

### Oxygen Consumption and Extracellular Acidification Rates by Seahorse XF Analyzer

1.25 × 10^5^ K562 cells were plated on a Seahorse plate in Seahorse XF DMEM media (Agilent) containing 25mM glucose and 4mM glutamine (ThermoFisher Scientific). Oxygen consumption and extracellular acidification rates were simultaneously recorded by a Seahorse XFe96 Analyzer (Agilent) using the mito stress test protocol, in which cells were sequentially perturbed by 2mM oligomycin, 1μM CCCP and 0.5mM antimycin (Sigma). Data were analyzed using the Seahorse Wave Desktop Software (Agilent). Data were not corrected for carbonic acid derived from respiratory CO_2_.

### Lactate Determination

Lacate secretion in the culture media was determined using a commercially available glycolysis cell-based assay kit (Cayman Chemical). An equal number of K562 cells expressing *GFP* or *UPP1*-FLAG were washed in PBS pre-incubated for 24h in no glucose DMEM media supplemented with 10% dialyzed fetal bovine serum (FBS, ThermoFisher Scientific), 100 U/mL penicillin/streptomycin (ThermoFisher Scientific) and 5mM of either glucose, galactose or uridine (all from Sigma) dissolved in water, or with an equal volume of water alone. Cells were then re-counted and seeded in fresh media of the same formulation and incubated for three additional hours. Cells were then spun down and lactate concentration was determined on the supernatants (spent media).

### Gene Ontology Analysis

Gene ontology analysis was performed using GOrilla with default settings and using a ranked gene list as input (*35*).

### Gene-Specific cDNA Cloning and Expression

cDNAs of interest were custom designed (Genewiz or IDT) and cloned into pWPI-Neo or pLV-lenti-puro using BamHI and SpeI (New-England Biolabs). cDNA sequences were:

#### *UPP1*-3xFLAG

GCTAGCGGATCCATGGCCGCCACCGGCGCCAACGCCGAGAAGGCCGAGAGCCACAACGACTGCCCCGTGAGGCTGCTGAACCCCAACATCGCCAAGATGAAGGAGGACATCCTGTACCACTTCAACCTGACCACCAGCAGGCACAACTTCCCCGCCCTGTTCGGCGACGTGAAGTTCGTGTGCGTGGGCGGCAGCCCCAGCAGGATGAAGGCCTTCATCAGGTGCGTGGGCGCCGAGCTGGGCCTGGACTGCCCCGGCAGGGACTACCCCAACATCTGCGCCGGCACCGACAGGTACGCCATGTACAAGGTGGGCCCCGTGCTGAGCGTGAGCCACGGCATGGGCATCCCCAGCATCAGCATCATGCTGCACGAGCTGATCAAGCTGCTGTACTACGCCAGGTGCAGCAACGTGACCATCATCAGGATCGGCACCAGCGGCGGCATCGGCCTGGAGCCCGGCACCGTGGTGATCACCGAGCAGGCCGTGGACACCTGCTTCAAGGCCGAGTTCGAGCAGATCGTGCTGGGCAAGAGGGTGATCAGGAAGACCGACCTGAACAAGAAGCTGGTGCAGGAGCTGCTGCTGTGCAGCGCCGAGCTGAGCGAGTTCACCACCGTGGTGGGCAACACCATGTGCACCCTGGACTTCTACGAGGGCCAGGGCAGGCTGGACGGCGCCCTGTGCAGCTACACCGAGAAGGACAAGCAGGCCTACCTGGAGGCCGCCTACGCCGCCGGCGTGAGGAACATCGAGATGGAGAGCAGCGTGTTCGCCGCCATGTGCAGCGCCTGCGGCCTGCAGGCCGCCGTGGTGTGCGTGACCCTGCTGAACAGGCTGGAGGGCGACCAGATCAGCAGCCCCAGGAACGTGCTGAGCGAGTACCAGCAGAGGCCCCAGAGGCTGGTGAGCTACTTCATCAAGAAGAAGCTGAGCAAGGCCgattataaagatcatgatggcgattataaagatcatgatattgattataaagatgatgatgataaataatagtgaGCGGCCGCACTAGTGAATTC

#### *UPP2*-3xFLAG

GCTAGCGGATCCATGGCCAGCGTGATCCCCGCCAGCAACAGGAGCATGAGGAGCGACAGGAACACCTACGTGGGCAAGAGGTTCGTGCACGTGAAGAACCCCTACCTGGACCTGATGGACGAGGACATCCTGTACCACCTGGACCTGGGCACCAAGACCCACAACCTGCCCGCCATGTTCGGCGACGTGAAGTTCGTGTGCGTGGGCGGCAGCCCCAACAGGATGAAGGCCTTCGCCCTGTTCATGCACAAGGAGCTGGGCTTCGAGGAGGCCGAGGAGGACATCAAGGACATCTGCGCCGGCACCGACAGGTACTGCATGTACAAGACCGGCCCCGTGCTGGCCATCAGCCACGGCATGGGCATCCCCAGCATCAGCATCATGCTGCACGAGCTGATCAAGCTGCTGCACCACGCCAGGTGCTGCGACGTGACCATCATCAGGATCGGCACCAGCGGCGGCATCGGCATCGCCCCCGGCACCGTGGTGATCACCGACATCGCCGTGGACAGCTTCTTCAAGCCCAGGTTCGAGCAGGTGATCCTGGACAACATCGTGACCAGGAGCACCGAGCTGGACAAGGAGCTGAGCGAGGAGCTGTTCAACTGCAGCAAGGAGATCCCCAACTTCCCCACCCTGGTGGGCCACACCATGTGCACCTACGACTTCTACGAGGGCCAGGGCAGGCTGGACGGCGCCCTGTGCAGCTTCAGCAGGGAGAAGAAGCTGGACTACCTGAAGAGGGCCTTCAAGGCCGGCGTGAGGAACATCGAGATGGAGAGCACCGTGTTCGCCGCCATGTGCGGCCTGTGCGGCCTGAAGGCCGCCGTGGTGTGCGTGACCCTGCTGGACAGGCTGGACTGCGACCAGATCAACCTGCCCCACGACGTGCTGGTGGAGTACCAGCAGAGGCCCCAGCTGCTGATCAGCAACTTCATCAGGAGGAGGCTGGGCCTGTGCGACgattataaagatcatgatggcgattataaagatcatgatattgattataaagatgatgatgataaataatagtgaGCGGCCGCACTAGTGAATTC

## Supplemental figures

**Figure S1:**
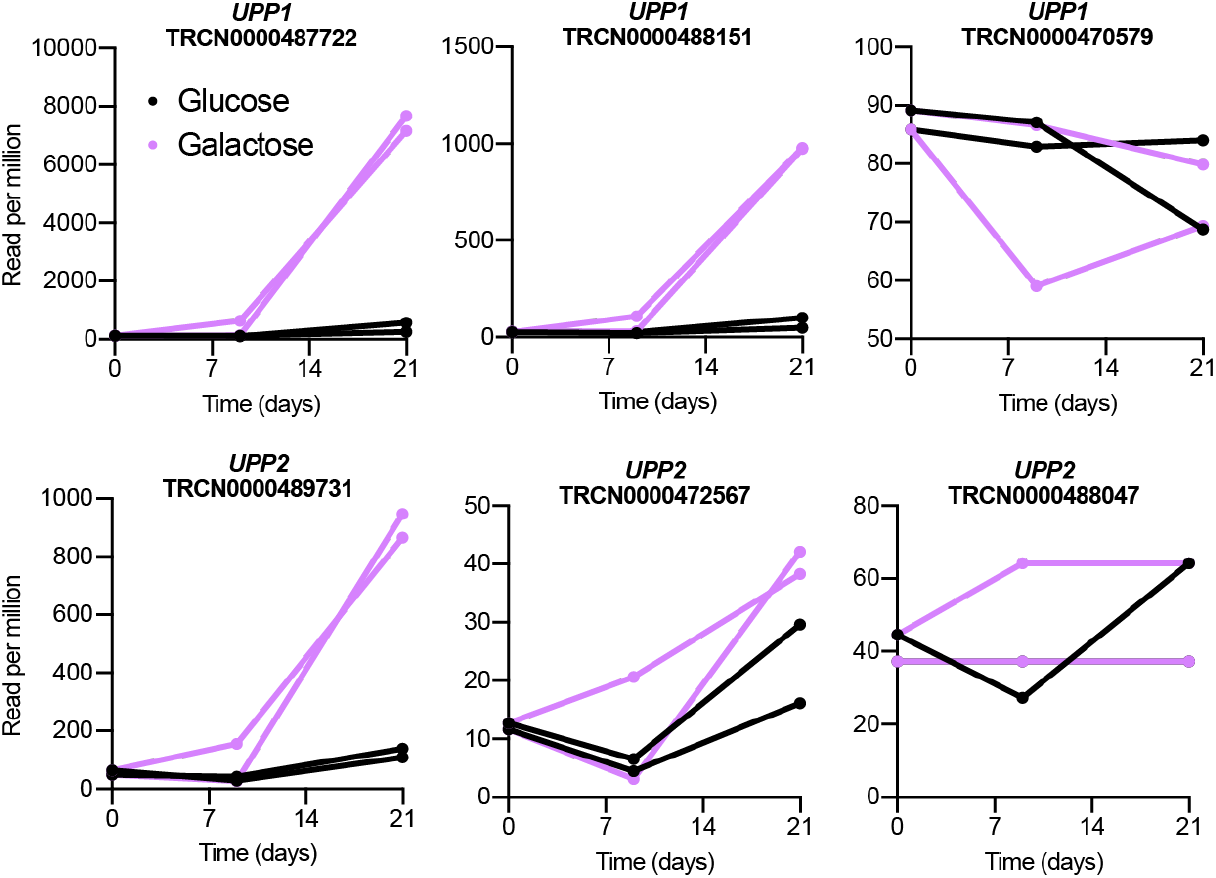
Additional Analysis of the ORF Screen. Representation of all six *UPP* ORFs as a function of time, expressed as read per million in the global population of glucose or galactose-grown cells (*n*=2).

**Figure S2:**
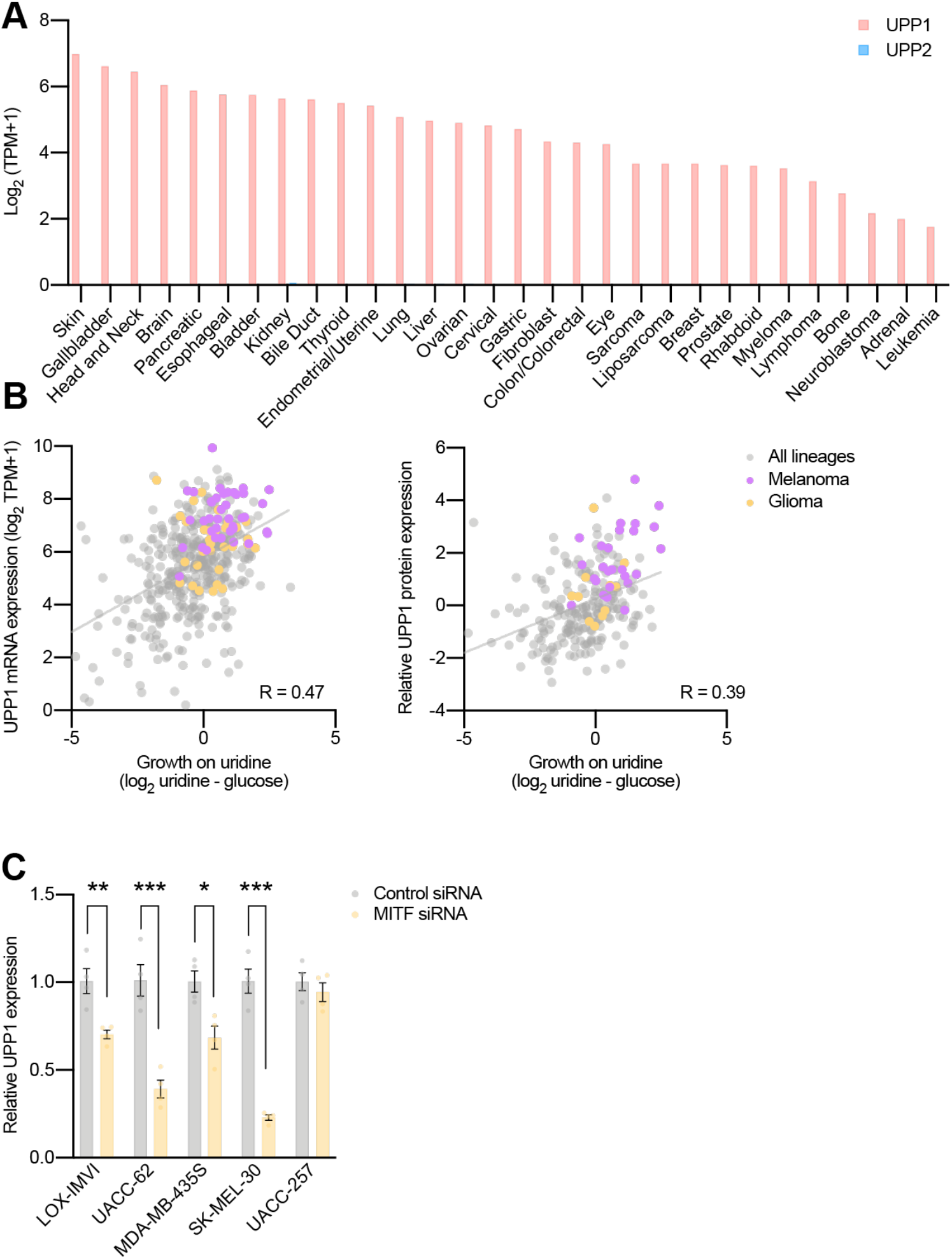
Additional Gene Expression and Correlation Analyses in the PRISM Collection. (A) *UPP1* and *UPP2* expression across the complete PRISM collection (*n*=30 lineages). (B) Correlation analysis comparing growth on uridine relative to glucose and the expression of *UPP1* transcripts and proteins across the 500 cancer cell lines from the PRISM screen, highlighting the melanoma and glioma lineages. (C) qPCR analysis of five melanoma cells after treatment with *MITF* siRNA. Data are shown as mean ±SEM with *p<0.05, **p<10^−2^ and ***p<10^−3^ t-test relative to the indicated control.

**Figure S3:**
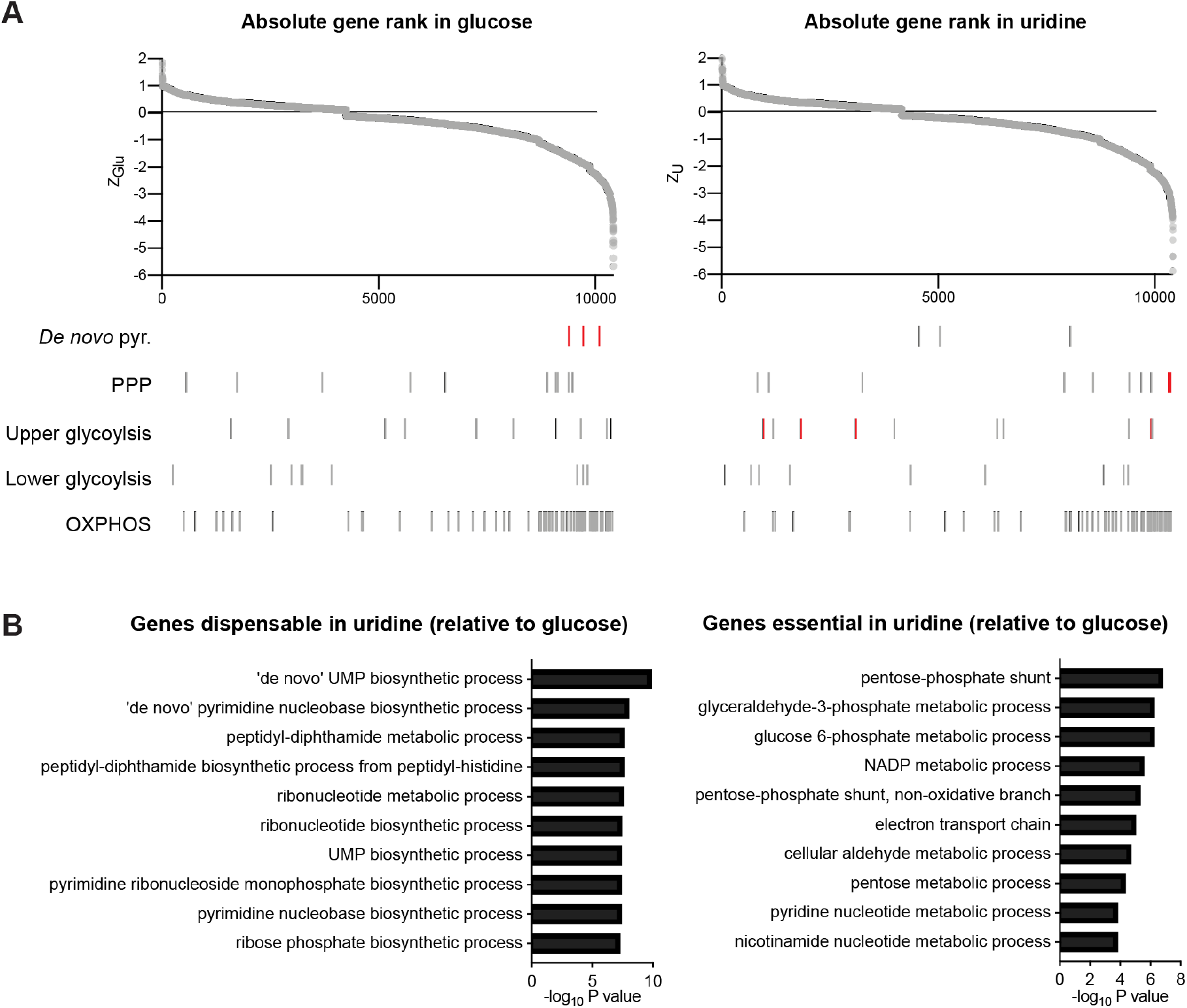
Genome-Wide CRISPR/Cas9 screen and Gene Ontology Analysis in Glucose and Uridine. (A) Genome-wide CRISPR/Cas9 screening reporting gene essentiality in glucose or in uridine. More negative Z scores indicated increased essentiality. Highlighted genes are discussed in the manuscript, and scores for all genes are reported in table S3. (B) Comparative gene ontology (GO) analysis highlighting the top 10 pathways in glucose *vs* uridine. GO was performed using GOrilla with default settings and using GO Process and a ranked gene list as input.

**Figure S4:**
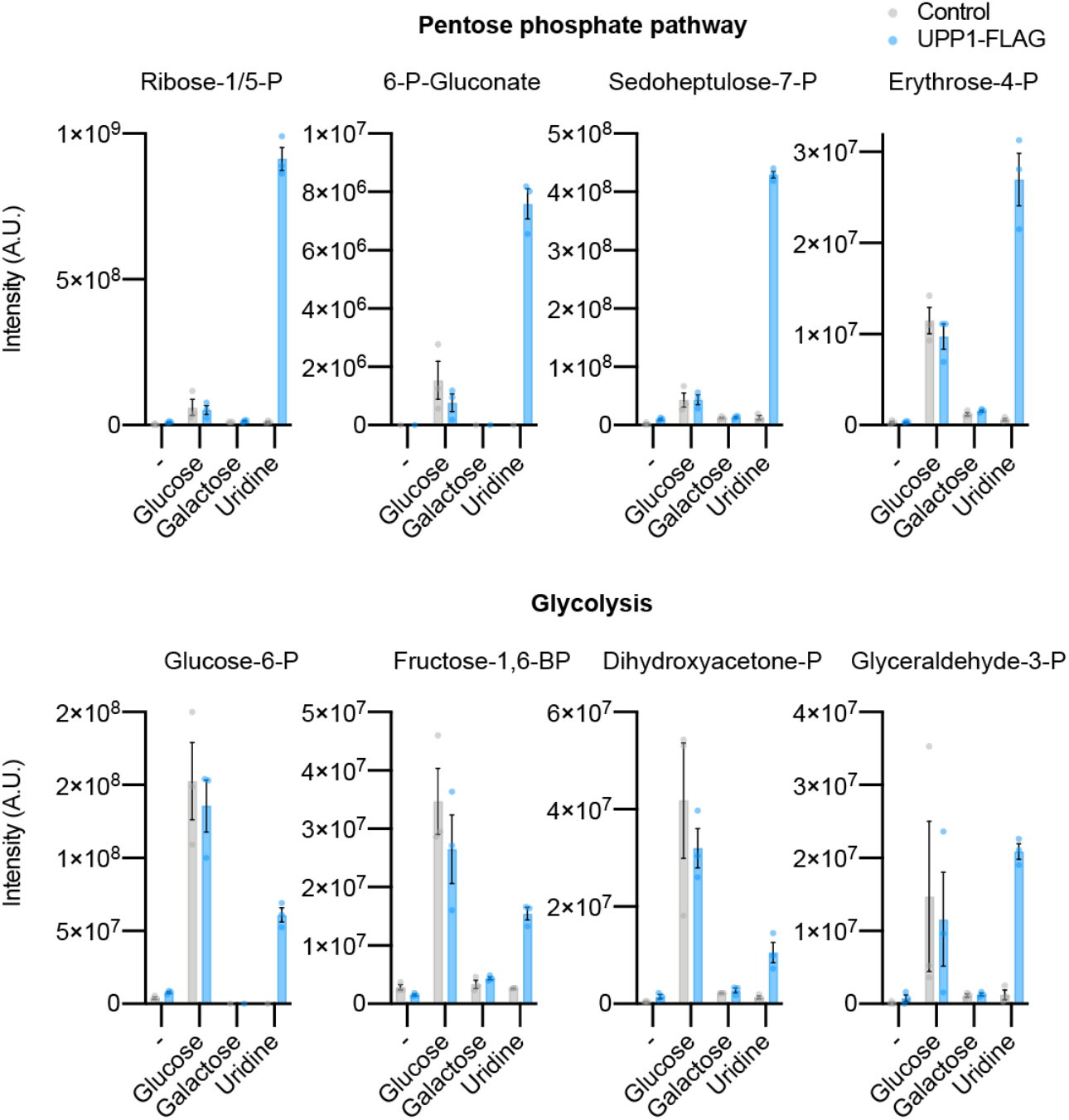
Steady-State Metabolomics Analysis of Uridine, Glucose and Galactose-Grown Cells. Steady-state abundance of representative intracellular metabolites from the pentose phosphate pathway (PPP) and glycolysis in sugar-free media complemented with 10mM of glucose, 10mM of galactose or 10 mM of uridine (*n*=3).

